# Continuous Theta Burst Stimulation to the Secondary Visual Cortex at 80% Active Motor Threshold Does Not Impair Central Vision in Humans During a Simple Detection Task

**DOI:** 10.1101/2021.04.11.439187

**Authors:** Carly A. Lasagna, Stephan F. Taylor, Taraz G. Lee, Saige Rutherford, Tristan Greathouse, Pan Gu, Ivy F. Tso

## Abstract

Continuous theta burst stimulation (cTBS) is a powerful form of repetitive transcranial magnetic stimulation capable of suppressing cortical excitability for up to 50 min. A growing number of studies have applied cTBS to the visual cortex in human subjects to investigate the neural dynamics of visual processing, but few have specifically examined its effects on central vision, which has crucial implications for safety and inference on downstream cognitive effects. The present study assessed the safety of offline, neuronavigated cTBS to V2 by examining its effects on central vision performance. In this single-blind, randomized sham-controlled, crossover study, 17 healthy adults received cTBS (at 80% active motor threshold) and sham to V2 1–2 weeks apart. Their central vision (≤8°) was tested at 1-min (T1) and again at 50-min (T50) post-stimulation. Effects of condition (cTBS vs. sham) and time (T1 vs. T50) on accuracy and reaction time were examined using Bayes factor. Bayes factor results suggested that cTBS did not impair stimulus detection over the entire central visual field nor subfields at T1 or T50. Our results offer the first explicit evidence supporting that cTBS applied to V2 does not create blind spots in the central visual field in humans during a simple detection task. Any subtler changes to vision and downstream visual perception should be investigated in future studies.

## INTRODUCTION

Transcranial magnetic stimulation (TMS) is a powerful, non-invasive technique for modulating cortical activity. When multiple pulses are delivered in patterned succession, as occurs in repetitive TMS (rTMS), it can excite or inhibit cortical activity within focal regions for sustained periods (Eldaief et al., 2011). In the field of cognitive neuroscience, low-frequency rTMS protocols (e.g., 1 Hz) – capable of eliciting long-term inhibition in the human cortex (Klomjai et al., 2015) – have become a popular non-invasive tool for elucidating the neural mechanisms of mental processes through induction of “virtual lesions.” In 2005, a patterned form of rTMS – continuous theta burst stimulation (cTBS), comprised of three pulses of stimulation at 50 Hz delivered every 200 ms for 40 s, for a total of 600 pulses (Huang et al., 2005) – has gained attention because this specific pattern is capable of suppressing activity for longer periods (up to 50-min) with briefer administration times than more non-patterned stimulation (40 s vs. 10–20 min) (Huang and Rothwell, 2004; Huang et al., 2005; Nyffeler et al., 2006; Thut and Pascual-Leone, 2010). As such, cTBS provides a quick, non-invasive, and well-tolerated (Huang et al., 2005) means to test the causal roles of specific brain regions in functional abnormalities observed in traumatic brain injuries or neuropsychiatric disorders.

The inhibitory effects of cTBS have been well-studied in motor cortices (Huang et al., 2005). Recent evidence suggesting that cTBS similarly suppresses cortical excitability when applied to *visual cortex* (Franca et al., 2006; Brückner and Kammer, 2016) has prompted studies to use cTBS to advance our understanding of brain dynamics underlying visual processing. For example, applications of cTBS to late-stage visual areas such as V5 have shown modulation of higher-level (global motion) processing without impairing lower-level (local motion) perception (Cai et al., 2014). However, when administering cTBS on early visual areas (i.e., V1–V3) to examine downstream effects, one critical challenge arises. Studies on cats have shown that brief TMS pulse trains (1–8 Hz for 1–4 s) to early occipital regions can generate visual field defects or scotoma for up to 10-min (Allen et al., 2007). Given cross-species similarities in the organization of these early areas (McKeefry et al., 2009), it is possible that cTBS applied to early visual areas in humans can also impair vision for extended periods. Without confirming the integrity of vision following stimulation, altered performance on visual tasks (Bertini et al., 2010; Rahnev et al., 2013; Cai et al., 2014; Chiou and Lambon Ralph, 2016; Chen et al., 2020) after cTBS to visual cortices could be merely due to blind spots, rather than the hypothesized mechanism of reduced feedforward activity from early visual areas to later processing areas. Impaired vision not only threatens the internal validity of the findings, but also constitutes a safety issue for human subjects.

Relatively few studies applying cTBS to early visual areas in humans have examined its effects on vision. Among those that have, many have focused on more peripheral locations in the visual field. From both the safety and scientific integrity perspectives, blind spots to the central visual field would be more problematic and thus warrant more attention for investigation. Additionally, findings from these studies have been inconsistent – reporting impairments (Rahnev et al., 2013; Cai et al., 2014; Chiou and Lambon Ralph, 2016; Chen et al., 2020), improvements (Waterston and Pack, 2010; Clavagnier et al., 2013), and non-significant changes (Waterston and Pack, 2010; Brückner and Kammer, 2014; Kaderali et al., 2015; Abuleil et al., 2021) to vision. This is likely due to variability in methods, including differences in stimulation intensity, performance measurement, target localization, and task demands. In many cases, these results were also based on extremely small samples (many had *N <* 10) (e.g., Waterston and Pack, 2010; Brückner and Kammer, 2014; Kaderali et al., 2015; Abuleil et al., 2021), and therefore negative findings could reflect a failure to reject the null hypothesis due to limited power rather than a true absence of stimulation effect on vision. As a result, it is unclear whether cTBS can be safely applied to early visual areas without inducing transient blind spots in central vision.

The present study assessed the effects of cTBS to an early visual area, V2, on central vision under a “typical” stimulation intensity level used in the majority of cTBS studies, 80% active motor threshold (Turi et al., 2021). We used a single-blind crossover design, in which participants received both conditions (randomized to receive either cTBS first or sham first) and underwent central vision testing at 1-min and 50-min post-stimulation. V2 was targeted because it is one of the earliest visual processing regions and has strong feedforward connections to higher-level processing areas in both the dorsal and ventral visual streams. We tested central vision (visual angle ≤8°) which is critical to performing most visual perception tasks and may be particularly susceptible to cTBS effects due to the size and depth of the cortical surface it occupies (Raz and Levin, 2014). We stimulated at 80% of active motor threshold (AMT) because the majority of cTBS studies use AMT to determine stimulation intensity, and of those, ∼90% utilize an intensity of 80% AMT (Turi et al., 2021). Thus, cTBS to V2 at 80% AMT can be considered a “standard” intensity for V2-cTBS and will allow us to determine whether central vision changes with safety and data integrity implications (i.e., causing blind spots) occur under a typical cTBS protocol.

Lesion studies in macaques show that V2 has unique functionality in visual detection dependent on task demands – V2 lesions impair complex but spare basic stimulus detection (Merigan et al., 1993). Here, “basic detection” requires accurate distinctions between coarsely discriminable stimuli (e.g., discriminate horizontal from vertical lines displayed on plain background), while “complex detection” places more demands on the visual system (e.g., judge orientation of shapes comprised of disconnected dots displayed against background distractors). Thus, we hypothesized that cTBS to V2 in humans would not impact detection accuracy on a basic visual task. Given previous reports of slowed visual detection following cTBS to early occipital targets (Fiori et al., 2015), we hypothesized that reaction time (RT) would increase after cTBS.

## METHODS

### Participants

Participants were *N* = 17 healthy adults (six females; age 24.8 ± 8.6; education 16.5 ± 2.3 years). Inclusion criteria consisted of the following: ages 18–55, visual acuity equal to or better than 20/30 on a Snellen chart, intact central/peripheral vision (test details below), no contraindication to TMS (see Rossi et al., 2009) or MRI, not taking psychotropic medication, no medical conditions with neurological sequelae (e.g., traumatic brain injury), no prior mental illness according to Structured Clinical Interview for DSM-IV-TR, non-patient version [SCID-NP (First et al., 2002)], and no substance abuse in past month (according to SCID-NP; corroborated by passing urine drug screenings in each visit).

### Procedure

The study protocol was approved by the Institutional Review Board at the University of Michigan Medical School and conducted in accordance with the Declaration of Helsinki. Prior to data collection, written informed consent was collected from all participants.

Participants completed three sessions: baseline, cTBS, and sham (**Figure 1**). Order for cTBS and sham sessions was counterbalanced across participants. At baseline, each participant completed a screening assessment, high-resolution T1-weighted (T1w) and T2-weighted (T2w) structural MRI scans, and a procedure to determine active motor threshold. Vision tests (central, acuity, peripheral) were administered to allow participants to become acclimated with the tasks and to confirm all had normal vision. TMS was delivered with a MagVenture MagPro X100 70 mm figure-8 shaped TMS coil (MCF-B70). In each stimulation session (cTBS, sham), central vision was tested ∼1 min after stimulation (“T1”) and then again at ∼50-min (“T50”) after completing a ∼40-min fMRI scan. Note that the fMRI scans/tasks were unrelated to the research question being addressed in this paper (whether cTBS applied to V2 at 80% AMT causes blind spots) and thus are outside the scope of the present paper and will be reported elsewhere. Beyond monitoring for common adverse events associated with TMS (e.g., headache, syncope), we also tested for additional vision-related adverse events following cTBS (and sham) that would pose a safety concern for participants. These “vision safety checks” involved a peripheral vision test and a visual acuity test that were administered at the end of each stimulation session, to ensure intact vision and safety before participants leave the laboratory.

**FIGURE 1.**
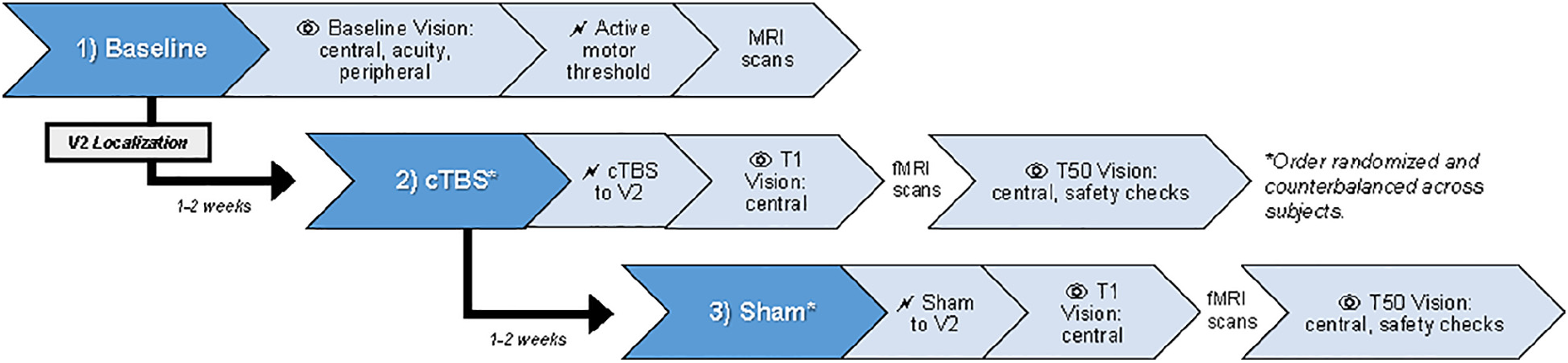
Overview of study procedures. Participants completed three sessions: baseline, cTBS, and sham-cTBS. Session order was randomized and counterbalanced across participants. T1 and T50, 1- and 50-min post-stimulation; cTBS, continuous theta burst stimulation; V2, secondary visual cortex. MRI scans were collected on Day 1 to guide target selection and TMS neuronavigation during subsequent visits. Note that the fMRI scans/tasks were unrelated to the research question being addressed in this paper (whether cTBS applied to V2 at 80% AMT causes blind spots) and thus are outside the scope of the present paper and will be reported elsewhere.

### Transcranial Magnetic Stimulation

#### Intensity

Stimulation intensity for the cTBS or sham was based on the participant’s active motor threshold (AMT), determined as the lowest intensity eliciting motor-evoked potentials of the first dorsal interosseous muscle (right hand; 20% maximum voluntary contraction) ≥100 µV on 5/10 trials. Mean raw AMT across participants was 36 ± 6% of maximum stimulator output (MSO).

#### Target Localization

Individual structural brain images were used to localize V2. High-resolution T1w and T2w anatomical images [256 × 256 FOV, 208 slices, 1 mm isotropic voxels, PROMO correction (White et al., 2010)] were acquired using a GE (MR750 DV25.0) 3T scanner and used to generate subject-specific masks of V2 and V1 (to facilitate determination of V2 boundaries). Details of image processing are provided in the **Supplementary Material**.

As shown in **Figure 2A**, V1 and V2 masks (right hemisphere) were superimposed on the native T1w volume in Brainsight software (Version 8; Rogue Research Inc., Montreal, QC, Canada). A target was placed in the center of a gyrus within the V2 cortex. Care was taken to avoid V1 and to ensure coil placement that minimized the scalp-to-target distance, as E-field strength reduces as a function of distance (Ilmoniemi et al., 1999). Across participants, the average distance from coil to target was 16.9 ± 3.7 mm. **Figure 2B** shows V2 targets for all subjects.

**FIGURE 2.**
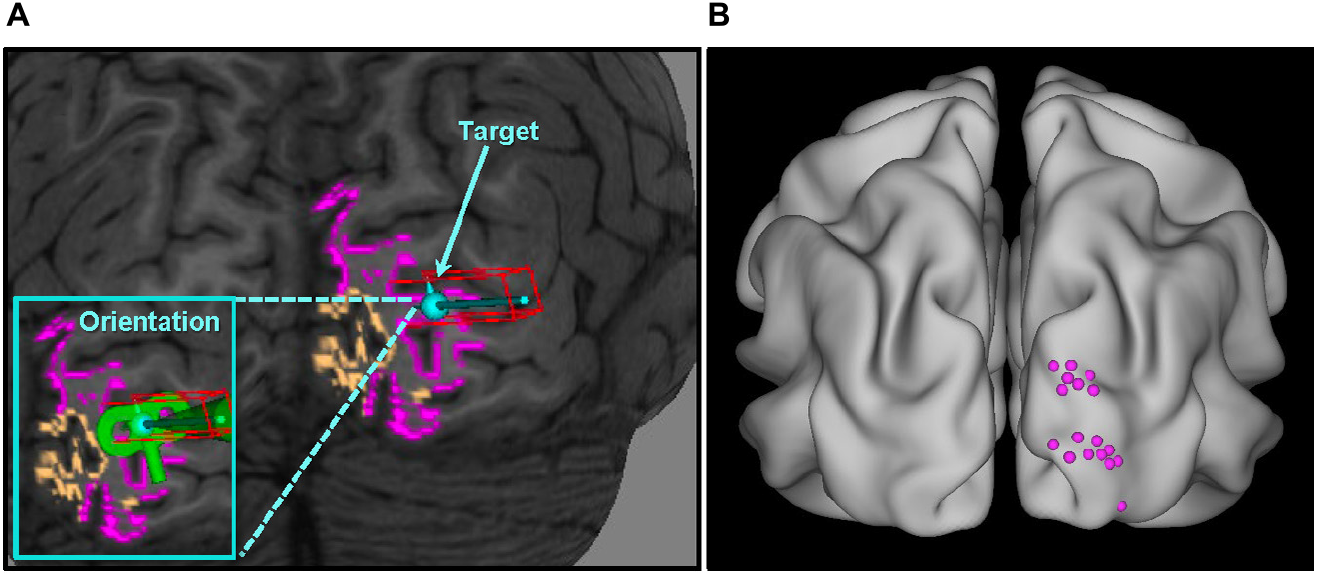
V2 localization overview. **(A)** Target selection for a single representative subject. Individualized overlays were superimposed on the T1 anatomical image (native space) – V1 colored orange, V2 colored pink – and used as guides in the V2 localization process; **(B)** V2 targets for all participants standardized in MNI space (average location: *x* = 21 ± 5, *y* = –95 ± 6, *z* = 19 ± 12; see **Supplementary Material** for individual subject coordinates). V1, primary visual cortex; V2, secondary visual cortex.

#### Continuous Theta Burst Stimulation

Continuous theta burst stimulation was delivered to V2 off-line at 80% of AMT with the coil oriented perpendicular to the gyrus of the target to optimize effects of the E-field (Janssen et al., 2015). Average stimulation intensity across participants was 29 ± 5% of MSO. cTBS parameters consisted of three 50 Hz pulse trains delivered every 200 ms continuously for 40 s, totaling 600 pulses (Huang et al., 2005). Brainsight Neuronavigation software with Polaris 3D Tracking (Version 8; Rogue Research Inc., Montreal, QC, Canada) utilizing T1w images enabled precise localization of target-centric subject/coil tracking during stimulation.

During stimulation, participants’ forehead was stabilized against a headrest to minimize movements. Given that research on state-dependent effects suggest that TMS to visual areas (V1– V4) maximally affects neuronal populations that are minimally active during stimulation (Silvanto et al., 2008), cTBS was administered while participants were blindfolded to control visual sensory input. For sham, the coil was rotated 90° so that the handle was perpendicular to the scalp.

### Vision Tests

#### Central Vision Task

A computerized task was used to examine binocular detection of visual stimuli across the central visual field (≤8°). Participants looked at a central fixation (size = 1°) while target stimuli (“1” or “2”; size = 0.75°) flashed briefly for 50 ms at different locations, one at a time, on the screen. Participants pressed the number key “1” or “2” to indicate what they saw and were given an indefinite period of time to respond. Following a response, a 200 ms interval elapsed before the next trial. Each trial was signaled by a 50 ms tone to minimize attention lapses. Modeled on Barendregt et al. (2014), targets were presented at 32 locations across the central visual field (**Figure 3**). These locations covered four visual angle eccentricities (2°, 4°, 6°, and 8°) and eight polarities rotated about a central fixation (22.5°, 67.5°, 112.5°, 157.5°, 202.5°, 247.5°, 292.5°, and 337.5°). Five presentations occurred at each of the 32 locations in randomized order, resulting in 160 trials and a task duration of ∼3-min. The task was presented in Psychtoolbox-3 (Brainard, 1997; Pelli, 1997; Kleiner et al., 2007) in MATLAB (R2019a) using a HP EliteBook laptop (Windows 10, 1920 × 1080 resolution, 33.5 cm × 17.5 cm display, 60 Hz refresh rate, 6-bit color depth) placed at an eye-level viewing distance of 55 cm. A headrest was used to minimize movements. Vision performance was evaluated by accuracy and RT for each stimulation condition (cTBS, sham) and administration time (T1, T50).

**FIGURE 3.**
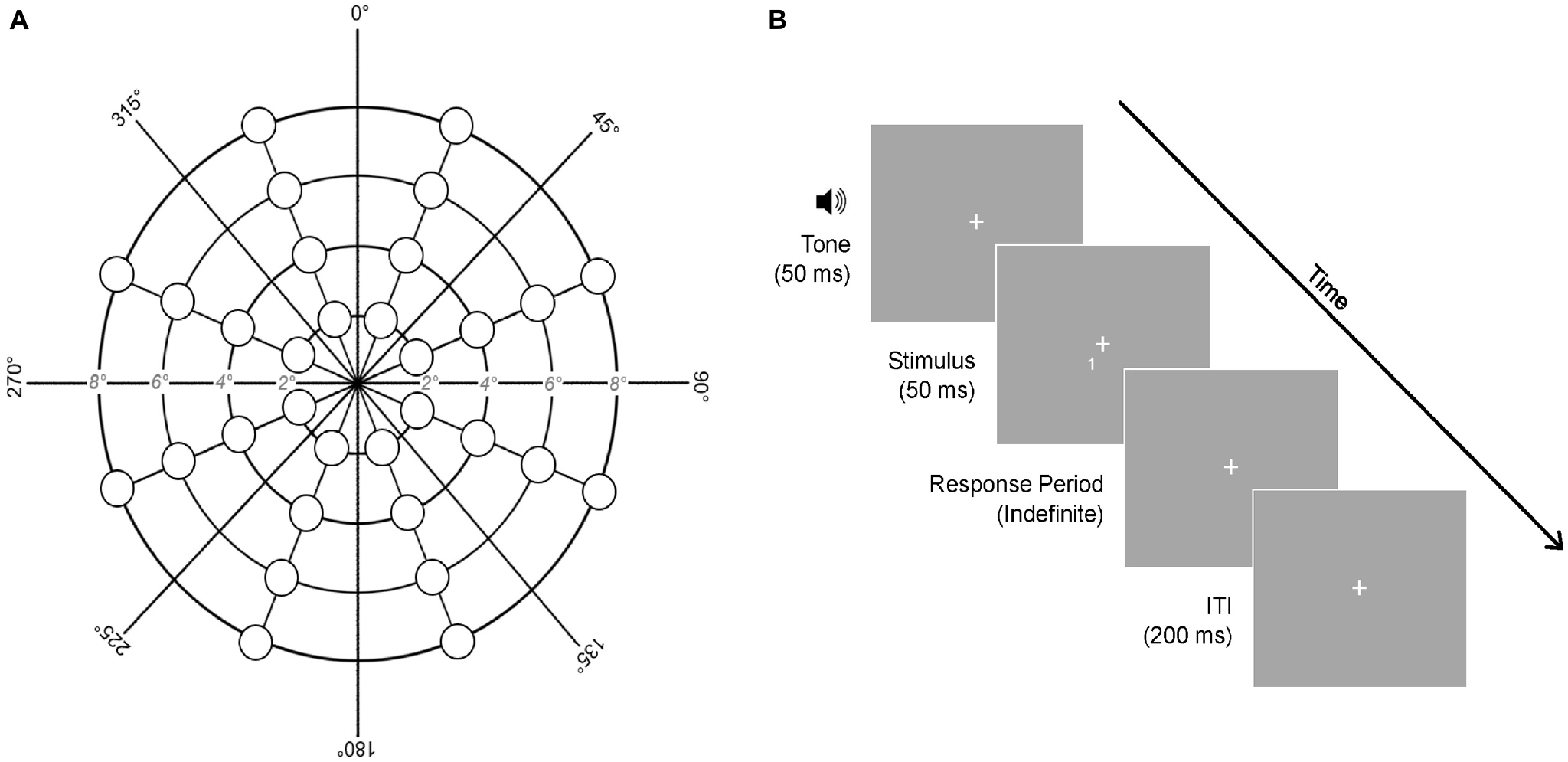
Central vision task. **(A)** Stimulus presentation locations. **(B)** Trial structure of the task. Each stimulus was preceded by a 50 ms tone, presented briefly for 50 ms, and followed by an indefinite response period. A 200 ms inter-trial interval (ITI) elapsed between trials. Five stimulus presentations occurred at each of the 32 locations shown, resulting in 160 trials. ms, milliseconds.

#### Vision Safety Checks

The integrity of peripheral vision was assessed monocularly using an experimenter-administered visual confrontation task (see **Supplementary Material** for description). Normal peripheral vision (requisite for participation) was defined as reliable identification on four trials in all quadrants of both eyes at baseline. Visual acuity was tested binocularly using a Snellen chart. Normal visual acuity (requisite for participation) was defined as 20/30 or better, with corrective lenses if needed. When re-tested after cTBS and sham stimulation sessions, any reductions in acuity or identification of stimuli in a quadrant were used to test for safety concerns related to stimulation.

### Statistical Analyses

The question of whether cTBS applied to an early visual area induces (temporary) blindness can be more meaningfully addressed by evaluating the *comparative* evidence for the null vs. alternative hypothesis. This can be achieved only by Bayesian analysis and not traditional null hypothesis significance testing. Therefore, we used a Bayesian model comparison approach to evaluate the relative strength of evidence for the null and alternative hypotheses (Quintana and Williams, 2018). We compared central vision performance following cTBS vs. sham by calculating the Bayes Factor (BF) – the ratio of Bayesian evidence for the alternative model (cTBS-related vision changes) to the null model (no cTBS-related vision changes). BF *<* 1 offers evidence for the null and BF *>* 1 offers evidence for the alternative. The strength of evidence is interpreted as follows (Jarosz and Wiley, 2014): evidence for null = 0.33–1 (anecdotal), 0.10–0.33 (substantial), 0.033–0.10 (strong), 0.01–0.033 (very strong), *<*0.01 (decisive); evidence for alternative = 1–3 (anecdotal), 3–10 (substantial), 10–30 (strong), 30–100 (very strong), *>*100 (decisive).

We used the “anovaBF” function in the “BayesFactor” (Morey and Rouder, 2015) R package to compute the BF of each possible alternative model with Stimulation (cTBS, sham) and Time (T1, T50) as possible fixed factors and subjects as a random effect, against the null model (subjects as random effect only) (Morey and Rouder, 2015). This was run separately for accuracy and RT as dependent variable. All outputs are provided in **Supplementary Material** and R analysis code is available at https://osf.io/3vxyn/.

## RESULTS

Continuous theta burst stimulation to V2 was well-tolerated and no participants reported common adverse side effects related to TMS (e.g., headache, syncope). Our vision safety checks revealed that no participants experienced reductions in visual acuity or impaired peripheral vision at the end of cTBS (or sham) visits. Accuracy and RT results are summarized in **Figure 4**.

**FIGURE 4.**
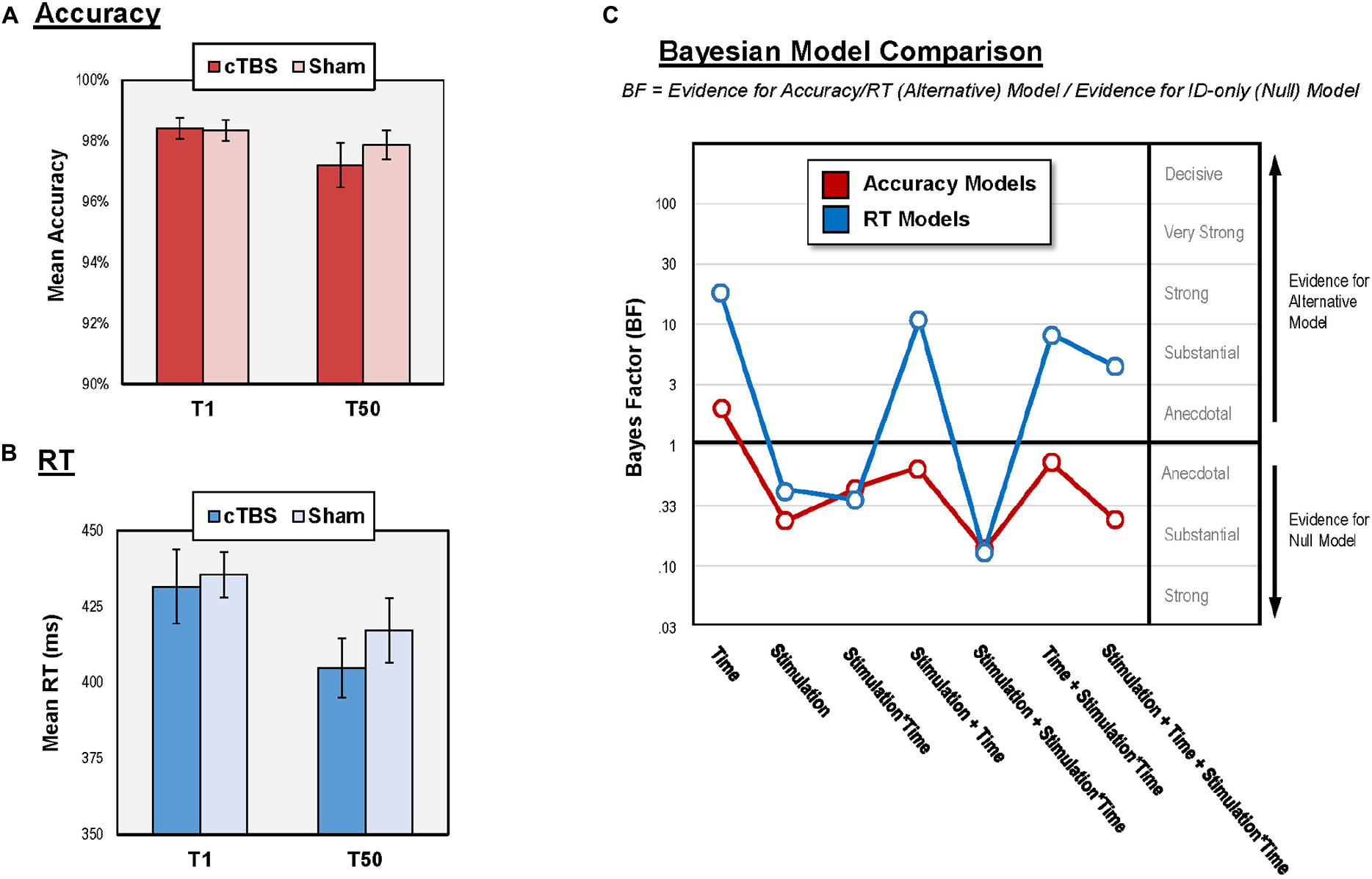
Results of central vision performance analysis. **(A)** Mean accuracy for each condition and administration time; **(B)** Mean RT for each condition and administration time; **(C)** Bayesian model comparison results for all models relative to ID-only model. BF *>* 1 suggest change in vision performance while BF *<* 1 suggest no change. RT, reaction time; cTBS, continuous theta burst stimulation; V2, secondary visual cortex; ms, milliseconds; T1 and T50, 1- and 50-min post-stimulation.

### Accuracy

Accuracy on the central vision task was high both after cTBS (T1 = 98.4 ± 5.8%; T50 = 97.2 ± 8.2%) and sham (T1 = 98.3 ± 5.8%; T50 = 97.9 ± 6.7%). Similarly, accuracy at all individual stimulus locations was also high (above 90% for all locations in both conditions). Vision performance for individual subjects is provided in **Supplementary Material**.

The null model had stronger evidence than all six models that contained Stimulation (as main effect or interaction with Time). Specifically, the evidence for the null model was “substantial” relative to three of these alternative models (BF = 0.31 to 0.14) and “anecdotal” relative to the other three alternative models (BF = 0.92 to 0.61). Together, the results suggest that cTBS to V2 did not impair accuracy of stimulus detection over the central visual field. The only alternative model with superior evidence than the null model was the one with Time as main effect (BF = 1.98, “anecdotal” evidence), indicating reduced accuracy at T50 relative to T1.

### RT

The null model had stronger evidence (“anecdotal” to “substantial” evidence; BF = 0.46 to 0.16) than three of the models containing Stimulation (as main effect or interaction with Time). The “winning” model with the strongest evidence contained Time as a main effect (BF = 24.91, “strong” evidence), suggesting learning effects for RT, marked by faster response at T50 relative to T1 in both stimulation conditions (**Figure 4B**). The other three alternative models containing Stimulation (as main effect or interaction with Time) had “substantial” to “strong” evidence relative the null (BF = 4.42 to 11.62). However, each of these alternative models also contained Time as a main effect. To confirm that these models won over the null model because of the Time effect, we performed comparisons showing that Stimulation did not affect RT *in addition to* Time. We compared the model with Time as a main effect against each of these three alternative models by dividing the BF of the former by that of the latter. In all cases, results favored the Time model, offering “anecdotal” to “substantial” evidence (BF = 2.14 to 5.63) supporting that Stimulation did not affect central vision RT in addition to Time. These results suggest that cTBS to V2 did not impact RT of detection over the central visual field immediately following or 50 min post-stimulation.

### Exploratory Analysis

We explored whether cTBS differentially impacted subareas of the visual field by considering the factors of Hemifield (ipsilateral vs. contralateral) and Vision Type [foveal (2°) vs. parafoveal (4– 8°)]. Neither Hemifield nor Vision Type was found to have effects on accuracy and RT (see **Supplementary Material**).

## DISCUSSION

The present study examined the effects of off-line, neuronavigated cTBS to V2 on central vision performance. Detection accuracy and RT over the central visual field were tested on a simple detection task following cTBS to V2 (at 1-min and 50-min post-stimulation) and compared to sham. We hypothesized that cTBS to V2 would not disrupt detection accuracy [based on results from lesion studies of the visual cortex (Merigan et al., 1993)] and would lead to slowed RT’s [based on previous TMS results (Fiori et al., 2015)].

As expected, cTBS to V2 did not affect the accuracy of central vision performance on a simple detection task. This held when hemifield and vision type were accounted for (see **Supplementary Material**), which helped rule out the possibility that cTBS to V2 differentially affected detection over a particular portion of the visual field. This extends on previous negative findings [obtained using null hypothesis significance testing (Brückner and Kammer, 2014; Kaderali et al., 2015)] related to discrimination accuracy following visual cortex cTBS. Our findings conflicted with a previous report of low-level visual processing impairment following cTBS to early visual areas (Rahnev et al., 2013), but this was likely due to the methodological differences. The Rahnev et al. (2013) study, along with the majority of previous studies on this topic (Waterston and Pack, 2010; Clavagnier et al., 2013; Nuruki et al., 2013; Rahnev et al., 2013; Brückner and Kammer, 2014; Kaderali et al., 2015), used the phosphene threshold (PT) hotspot as their “early visual” target, whereas we used an anatomical map of V2 (Glasser et al., 2016). Phosphene-based methods of localization can yield targets that vary widely between V1 through V3 (Schaeffner and Welchman, 2017). This can be problematic because, from lesion studies, we know that disruptions to different functional areas within “early occipital” regions have drastically different effects on visual processing [e.g., V1 lesions impair basic and complex visual processing, while V2 lesions impair complex processing only (Merigan et al., 1993)]. It is, therefore, possible that impairments reported in that study were caused by suppression of activity in V1 in some participants, which would be more likely to cause blind spots. In contrast, given our localization method, we were confident that V2 was stimulated in our study and thus our findings were specific to cTBS applied to V2 (see below for discussion of possible influences of intensity).

Contrary to hypotheses, cTBS to V2 did not impact RT of detection over the overall visual field nor specific subfields on a simple detection task. The discrepancy between this result and one previous study reporting slower RT following cTBS (Fiori et al., 2015) could be due to methodological differences. The task in the current study was relatively easy – a basic detection task with a fixed stimulus presentation duration (50 ms) – while Fiori et al. (2015) utilized staircase adaptation procedures to manipulate difficulty level, reducing stimuli viewing time in order to keep performance at 80% accuracy. The stimuli also varied in terms of complexity: we used coarsely discriminable targets while Fiori used more complex Gabor elements with varied orientations. Together, this placed more demands on participants in Fiori et al.’s (2015) study than those in the present study. This is important because, as discussed previously, lesions to V1 in macaques cause deficits on basic visual processing tasks, while lesions to V2 leave basic discrimination and contrast sensitivity unaffected (Merigan et al., 1993) but impair complex visual processing. Perhaps task demand/complexity is another reason why some have reported low-level visual impairments (Rahnev et al., 2013; Fiori et al., 2015) after early occipital cTBS, while our study and those using easier tasks reported no effects (Brückner and Kammer, 2014). Others have urged the importance of task complexity in TMS research (Waterston and Pack, 2010). Here we add that it is also important to carefully consider the specific visual area stimulated, because failure to consider both may have unforeseen interaction effects on vision performance.

Regardless of stimulation condition, time appeared to impact accuracy and RT, such that responses were less accurate but faster at T50 (relative to T1). It may be that participants were more comfortable with the task and thus were faster to respond (at a slight expense of accuracy).

Finally, one might question whether the stimulation intensity used in the present study was insufficient to adequately suppress V2 excitation. Although the intensity we used (80% AMT) was lower than others (i.e., using resting motor threshold, phosphene threshold, or a fixed percentage of maximum stimulator output), several studies have successfully modulated behavior, perception, and cognition using visual cortex cTBS at intensities similar to ours (van Nuenen et al., 2012; Beck et al., 2015; Adam et al., 2016). This suggests that the absence of cTBS effects on simple visual detection performance in this study was not likely due to insufficient stimulation power. While visual impairments may be possible at much higher intensities of cTBS to V2, our goal was *not* to demonstrate this *per se*. Rather, we sought to assess whether blind spots would occur under a standard intensity (80% AMT) (Rossi et al., 2009) most commonly adopted in cTBS research (Turi et al., 2021). In doing so, we demonstrated that cTBS delivered at standard intensities to V2 does not impair basic detection over the central visual field. However, this does not imply that cTBS to V2 at *any* intensities would leave vision intact. There is evidence that higher-intensity cTBS to early visual areas is capable of altering visual processing, but results are heterogeneous (Bertini et al., 2010; Waterston and Pack, 2010; Clavagnier et al., 2013; Chen et al., 2020).

These findings should be considered in light of several limitations. Our sample was modest and replications with larger sample sizes are needed. Additionally, the extant TMS literature (especially in the motor cortex) indicates that peak effects may not occur until ∼10–20 min after stimulation (Huang et al., 2005). Therefore, the testing points used here might not have captured possible disruptions associated with peak effects. Future work should use additional time intervals between 0- and 50- min following stimulation to assess for possible vision changes post-cTBS. This would be an important direction in future investigations to fully understand the time course of cTBS effects on central vision.

In summary, cTBS delivered at 80% AMT to V2 did not impair accuracy or RT of central vision detection during a simple detection task. Our findings provide further evidence consistent with previous reports that cTBS can be safely applied to V2 at standard intensities (80% AMT) and does not disrupt basic early visual detection needed to perform tasks tapping higher-level visual processing or cognition. Replications with larger samples are needed to provide more definitive conclusions about the safety of cTBS in early visual cortices and effects on vision and behavior.

## DATA AVAILABILITY STATEMENT

The raw data supporting the conclusions of this article will be made available by the authors, without undue reservation.

## ETHICS STATEMENT

The studies involving human participants were reviewed and approved by the University of Michigan Medical School IRBMED. The patients/participants provided their written informed consent to participate in this study.

## AUTHOR CONTRIBUTIONS

CL: methodology, software, investigation, data curation, formal analysis, visualization, and writing – original draft. ST: methodology, supervision, and writing – review and editing. TL: methodology, resources, supervision, and writing – review and editing. SR: methodology, software, and writing – review and editing. TG: software, formal analysis, visualization, and writing – review and editing. PG: investigation, data curation, visualization, and writing – review and editing. IT: conceptualization, methodology, funding acquisition, project administration, supervision, formal analysis, and writing – review and editing. All authors contributed to the article and approved the submitted version.

## ACKNOWLEDGMENTS

Portions of these results were presented at the 31st Annual Albert J. Silverman Research Conference, University of Michigan.

## SUPPLEMENTARY MATERIAL

The Supplementary Material for this article can be found online at: https://www.frontiersin.org/articles/10.3389/fnhum.2021.709275/full#supplementary-material

## Publisher’s Note

All claims expressed in this article are solely those of the authors and do not necessarily represent those of their affiliated organizations, or those of the publisher, the editors and the reviewers. Any product that may be evaluated in this article, or claim that may be made by its manufacturer, is not guaranteed or endorsed by the publisher.

*Copyright © 2021 Lasagna, Taylor, Lee, Rutherford, Greathouse, Gu and Tso. This is an open-access article distributed under the terms of the Creative Commons [Attribution License (CC BY)](https://creativecommons.org/licenses/by/4.0/). The use, distribution or reproduction in other forums is permitted, provided the original author(s) and the copyright owner(s) are credited and that the original publication in this journal is cited, in accordance with accepted academic practice. No use, distribution or reproduction is permitted which does not comply with these terms.*

