## Supplement for "Continuous Theta Burst Stimulation to the Secondary Visual Cortex at 80% Active Motor Threshold Does Not Impair Central Vision in Humans During a Simple Detection Task"

### *Supplementary Material*

#### 1 Supplementary information

##### 1.1 Visual confrontation task description

During the visual confrontation task, the participant sat one meter from the examiner at eye-level, with one eye covered, and fixated on the examiner's nose. The examiner established the limits of peripheral vision in all quadrants by moving their wagging finger from outside the visual field, towards a central position until the participant confirmed visualization. At the point of detection, the examiner randomly flashed 1 or 2 fingers on their right or left hand, to which the participant reported the side and number that appeared (e.g. "2 on my left"). This process was repeated until the participant achieved reliable identification on 4 trials for all quadrants, independently for each eye.

##### 1.2 MRI image processing

Images were processed offline in Freesurfer image analysis suite (version 6.0; http://surfer.nmr.mgh.harvard.edu/), which was used for cortical reconstruction and volumetric segmentation, using both T1w and T2w volumes (T2w images improve the quality of the pial surface reconstruction and thus the accuracy of the cortical surface map). Technical details of these procedures are described elsewhere (Bruce Fischl et al. 2002; Dale, Fischl, and Sereno 1999; B Fischl and Dale 2000; Desikan et al. 2006). Briefly, we conducted motion correction of volumetric images, removal of non-brain tissue, automated Talairach transformation, segmentation of subcortical white matter and deep gray matter volumetric structures (including hippocampus, amygdala, caudate, putamen, ventricles), intensity normalization, tessellation of gray matter white matter boundary, automated topology correction, and surface deformation. Individual V1 and V2 masks were generated during the surface reconstruction (recon-all) process.

##### 1.3 Exploratory analysis of V2-cTBS effects across the visual field

Exploratory analyses also assessed whether cTBS differentially impacted certain areas of the visual field. Accuracy and RT scores were computed for stimuli presented to each 'hemifield' (ipsilateral vs. contralateral to stimulation site) and 'vision type' (foveal [2°] vs. parafoveal [4-8°] visual fields). AnovaBF in R Studio was used to compute an alternative model containing Stimulation as an interaction with Hemifield (as well as random subjects effects), which was compared against a null model containing only random subjects effects. This process was also completed for an alternative model containing Stimulation as an interaction with Vision Type. All exploratory analyses were conducted separately for accuracy and RT data at T1 and T50 (i.e., models for accuracy-T1, accuracy-T50, RT-T1, RT-T50 were analyzed individually).

R code and outputs for exploratory accuracy and RT analyses are provided in Supplementary Code 3.1 and 3.2, respectively.

For accuracy, at T1 and T50, the alternative model that contained Stimulation as an interaction with Hemifield had weaker evidence than the null model (both BF = 0.33; 'anecdotal'). The same was true of the alternative model containing the interaction between Stimulation and Vision Type, which also had weaker evidence than the null model at T1 (BF = 0.33, 'anecdotal' evidence) and T50 (BF = 0.86, 'anecdotal' evidence). Together, these results suggest that cTBS did not impair central vision accuracy in a particular region of the visual field.

For RT, the alternative model containing Stimulation as an interaction with Vision Type had weaker evidence than the null model at both T1 (BF = 0.52; ‘anecdotal’) and T50 (BF = 0.34; ‘anecdotal’). The same was true of the alternative model containing the interaction between Stimulation and Vision Type, which also had weaker evidence than the null model at T1 (BF = 0.34, ‘anecdotal’ evidence) and T50 (BF = 0.50, ‘anecdotal’ evidence). Together, this indicates that cTBS did not slow central vision RT in a particular region of the visual field.

### 2 Supplementary figures and tables

#### 2.1 Individual V2 targets for *N* = 17 subjects standardized in MNI space.

| *Target MNI Coordinates* | | | |
|---|---|---|---|
| **ID** | **x** | **y** | **z** |
| 1 | 27 | -96 | 10 |
| 2 | 30 | -93 | -8 |
| 3 | 28 | -94 | 6 |
| 4 | 17 | -83 | 45 |
| 5 | 18 | -92 | 28 |
| 6 | 28 | -97 | 8 |
| 7 | 21 | -99 | 13 |
| 8 | 19 | -93 | 25 |
| 9 | 15 | -89 | 29 |
| 10 | 18 | -97 | 10 |
| 11 | 17 | -113 | 31 |
| 12 | 23 | -98 | 10 |
| 13 | 19 | -91 | 26 |
| 14 | 24 | -92 | 26 |
| 15 | 14 | -99 | 14 |
| 16 | 14 | -94 | 27 |
| 17 | 23 | -90 | 24 |
| **Mean** | **21** | **-95** | **19** |
| **SD** | **5** | **6** | **12** |

***Targets on Canonical Brain***

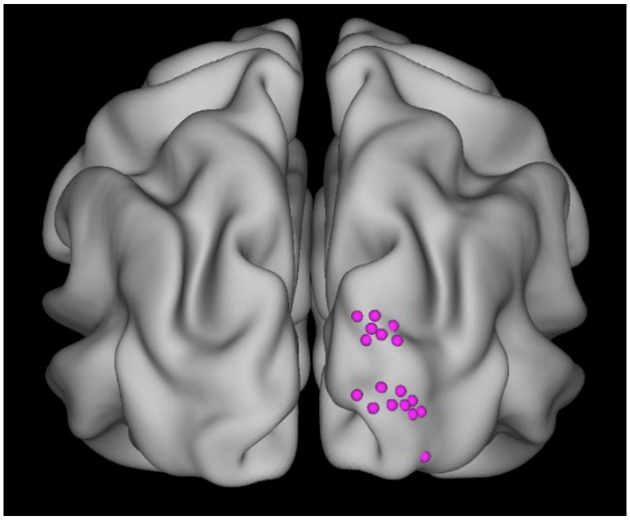

#### 2.1 Individual central visual performance

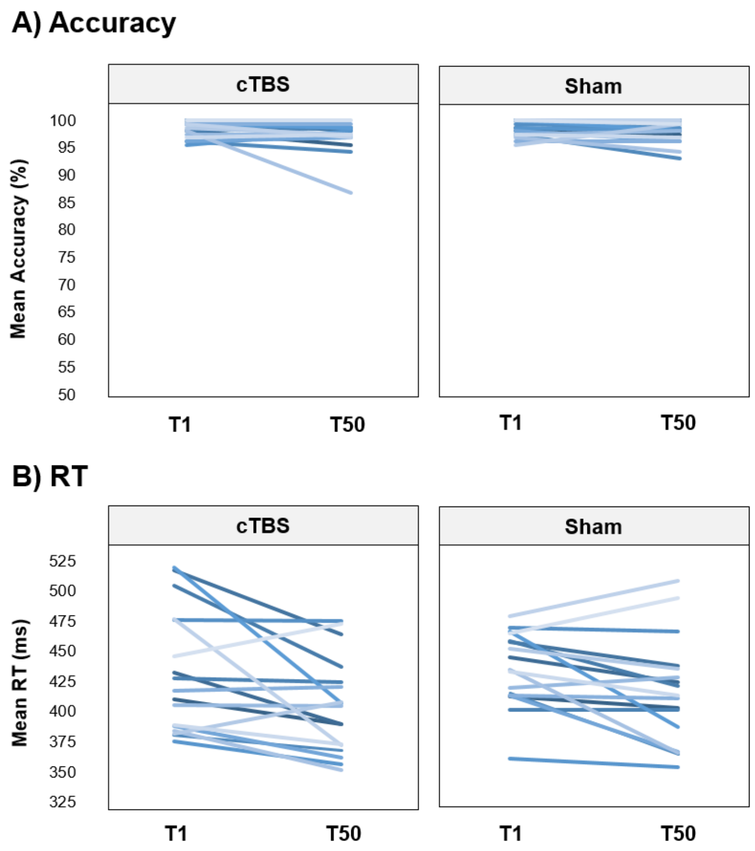


**Central vision performance for *N* = 17 individual participants.** Values shown represent mean accuracy and RT during central vision testing at T1 (1-minute post stimulation) and T50 (50-minutes post-stimulation) after cTBS and sham to V2. *Note:* cTBS = continuous theta burst stimulation; RT = reaction time.

#### 3. Supplementary code and outputs

##### 3.1 Accuracy Bayes factor analyses (complete R code available at: https://osf.io/3vxyn/)

<u>CENTRAL ACCURACY ANALYSES</u>

**Analysis 1:** Generate all accuracy models. Include fixed factors of STIM and TIME and additive random subjects effects (ID). Calculate BF of all models compared to ID-only model.

| | BF Analysis Output | | Evidence Interpretation | |
|---|---|---|---|---|
| | **BF** | **Error Interval** | **Favors** | **Magnitude** |
| [1] stim + id | 0.309 | [0.306 0.313] | Null | Substantial |
| [2] time + id | 1.983 | [1.939 2.027] | Alternative | Anecdotal |
| [3] stim*time + id | 0.461 | [0.456 0.467] | Null | Anecdotal |
| [4] stim + time + id | 0.612 | [0.595 0.628] | Null | Anecdotal |
| [5] stim + stim*time + id | 0.144 | [0.141 0.147] | Null | Substantial |
| [6] time + stim*time + id | 0.918 | [0.894 0.942] | Null | Anecdotal |
| [7] stim + time + stim*time + id | 0.286 | [0.28 0.292] | Null | Substantial |

**EXPLORATORY ACCURACY ANALYSES**

**Analysis 2:** Check for possible effects of VISION TYPE. Generate exploratory acc models separately for T1. Include fixed factors of STIM and VISION_TYPE (foveal/parafoeval visual field) and additive random subjects effects (ID). Then do the same for T50.

| | BF Analysis Output | | Evidence Interpretation | |
|---|---|---|---|---|
| | **BF** | **Error Interval** | **Favors** | **Magnitude** |
| [1] stim_t1*vision_type_t1 + id_t1 | 0.326 | [0.319 0.334] | Null | Substantial |

| | BF Analysis Output | | Evidence Interpretation | |
|---|---|---|---|---|
| | **BF** | **Error Interval** | **Favors** | **Magnitude** |
| [1] stim_t50*vision_type_t50 + id_t50 | 0.331 | [0.329 0.334] | Null | Substantial |

**Analysis 3:** Check for possible effects of stimulus HEMIFIELD. Generate exploratory acc models separately for T1. Include fixed factors of STIM and HEMIFIELD (contralateral/ipsilateral to stimulation site) and additive random subjects effects (ID). Then do the same for T50.

| | BF Analysis Output | | Evidence Interpretation | |
|---|---|---|---|---|
| | **BF** | **Error Interval** | **Favors** | **Magnitude** |
| [1] stim_t1*hemifield_t1 + id_t1 | 0.327 | [0.323 0.33] | Null | Substantial |

| | BF Analysis Output | | Evidence Interpretation | |
|---|---|---|---|---|
| | **BF** | **Error Interval** | **Favors** | **Magnitude** |
| [1] stim_t50*hemifield_t50 + id_t50 | 0.88 | [0.853 0.898] | Null | Anecdotal |

#### 3.2 **RT Bayes factor analyses** (complete R code available at: https://osf.io/3vxyn/)

**CENTRAL RT ANALYSES**

**Analysis 4:** Generate all RT models. Include fixed factors of STIM and TIME and additive random subjects effects (ID). Calculate BF of all models compared to ID-only model.

| | BF Analysis Output | | Evidence Interpretation | |
|---|---|---|---|---|
| | **BF** | **Error Interval** | **Favors** | **Magnitude** |
| [1] stim + id | 0.461 | [0.43 0.492] | Null | Anecdotal |
| [2] time + id | 24.909 | [24.603 25.216] | Alternative | Strong |
| [3] stim*time + id | 0.367 | [0.364 0.371] | Null | Anecdotal |
| [4] stim + time + id | 11.618 | [11.355 11.882] | Alternative | Strong |
| [5] stim + stim*time + id | 0.161 | [0.158 0.164] | Null | Substantial |
| [6] time + stim*time + id | 9.111 | [8.948 9.274] | Alternative | Substantial |
| [7] stim + time + stim*time + id | 4.421 | [4.299 4.544] | Alternative | Substantial |

**EXPLORATORY RT ANALYSES**

**Analysis 5:** Check for possible effects of VISION TYPE. Generate exploratory rt models separately for T1. Include fixed factors of STIM and VISION_TYPE (foveal/parafoeval visual field) and additive random subjects effects (ID). Then do the same for T50.

| | BF Analysis Output | | Evidence Interpretation | |
|---|---|---|---|---|
| | **BF** | **Error Interval** | **Favors** | **Magnitude** |
| [1] stim_t1*vision_type_t1 + id_t1 | 0.516 | [0.395 0.637] | Null | Anecdotal |

| | BF Analysis Output | | Evidence Interpretation | |
|---|---|---|---|---|
| | BF | Error Interval | Favors | Magnitude |
| [1] stim_t50*vision_type_t50 + id_t50 | 0.340 | [0.335 0.345] | Null | Anecdotal |

**Analysis 6:** Check for possible effects of stimulus HEMIFIELD. Generate exploratory rt models separately for T1. Include fixed factors of STIM and HEMIFIELD (contralateral/ipsilateral to stimulation site) and additive random subjects effects (ID). Then do the same for T50.

| | BF Analysis Output | | Evidence Interpretation | |
|---|---|---|---|---|
| | BF | Error Interval | Favors | Magnitude |
| [1] stim_t1*hemifield_t1 + id_t1 | 0.344 | [0.336 0.352] | Null | Anecdotal |

| | BF Analysis Output | | Evidence Interpretation | |
|---|---|---|---|---|
| | BF | Error Interval | Favors | Magnitude |
| [1] stim_t50*hemifield_t50 + id_t50 | 0.500 | [0.487 0.513] | Null | Anecdotal |
